# Mycobacterial DNA-binding protein 1 (MDP1) induces RNA condensation as revealed by high-speed AFM

**DOI:** 10.1101/2025.08.03.668364

**Authors:** Nadia Shoukat, Akihito Nishiyama, Kaho Nakamoto, Yuna Goto, Phuong Doan N. Nguyen, Hikaru Ichida, Kenichi Umeda, Sohkichi Matsumoto, Noriyuki Kodera

## Abstract

Mycobacterial DNA-binding protein 1 (MDP1) is a histone-like protein in *Mycobacterium tuberculosis* (Mtb) that contributes to genome organization and dormancy adaptation. MDP1 consists of a structured HU-like region (HUR), and a post-translational modifications (PTMs)-enriched intrinsically disordered region (IDR), that regulate its function. While its role in DNA condensation is well established, how MDP1 interacts with RNA remains unclear, despite growing recognition of transcriptional regulation during dormancy. Here, by using total RNA from *E. coli* as a model substrate, we investigated whether MDP1 induces RNA condensation. Our high-speed AFM and optical microscopy analysis shows that native MDP1 from Mtb, rich in PTMs, forms globular RNA condensates. In contrast, MDP1 expressed in *E. coli*, which lacks PTMs, induces chain-like RNA condensates. Domain-specific analysis revealed that the IDR, the synergy between the IDR and HUR, and PTMs are essential for MDP1-induced RNA condensation. These findings suggest that MDP1 mediates RNA organization during dormancy.

## TEXT

Tuberculosis (TB), caused by *Mycobacterium tuberculosis* (Mtb), remains one of the deadliest infectious diseases globally. According to the World Health Organization (WHO), an estimated 10.8 million people developed TB in 2023, and 1.25 million died from the disease. After being overtaken by COVID-19 for three consecutive years, TB has resumed its status as the leading cause of mortality due to a single infectious pathogen^1^.

A key factor in the persistence of TB is the ability of Mtb to enter a dormant state, during which the bacterium becomes metabolically inactive and highly resistant to antibiotics and immune clearance^2–7^. Dormancy is accompanied by a profound metabolic slowdown, extensive reorganization of nucleic acids, and transcriptional reprogramming, enabling Mtb to survive within the host for decades^2,3,8,9^. This condition known as latent TB infection affects nearly one-quarter of the global population^3,10^. Understanding the molecular mechanisms underlying dormancy is therefore crucial for developing effective therapies, especially in light of the growing burden of drug-resistant TB, which accounted for ∼0.4 million cases of multidrug-resistant TB in 2023^11^.

A key player in Mtb dormancy is Mycobacterial DNA-binding protein 1 (MDP1), also known as Rv2986c, MtHU, HU (histone like protein)-homolog or hupB^4^. MDP1, is a nucleoid-associated, histone-like protein composed of 214 amino acids with an approximate molecular weight of 21 kDa^12^. It contains a conserved N-terminal domain (residues 1–99) that is homologous to bacterial HU proteins, referred to here as the HU-like region (HUR), and a C-terminal intrinsically disordered region (IDR; residues 100– 214) enriched in post-translational modifications (PTMs)^13,14^. These PTMs, which include mono-, di-, and trimethylation, acetylation and succinylation, predominantly occur within the IDR and have been shown to modulate the structural and function of MDP1^13,15^.

The primary role of MDP1 in DNA compaction has been well-documented^16–19^. Our recent study using high-speed atomic force microscopy (HS-AFM), a powerful technique capable of nano-scale video imaging of macromolecules in action^20–22^, provided a detailed molecular-level view of this process in which MDP1 cross-links two DNA duplexes using IDR like a double-sided tape^3^. This nanoscopic observation offers new mechanistic understanding of MDP1-mediated DNA organization^16^.

In addition to DNA compaction, transcriptomic studies indicate that RNA is also selectively regulated during dormancy, raising the possibility that RNA organization may contribute to stress adaptation^8,9,14,23^. While we have reported the binding of MDP1 to RNA *in-vitro*^23^, the structural and regulatory consequences of this interaction, particularly in the context of RNA condensation and its potential PTM dependence remain unexplored, leaving a significant gap in our understanding of Mtb dormancy mechanisms.

Here, we address this question using a combination of HS-AFM and optical microscopy (OM). In our previous study, we demonstrated that MDP1 binds to MS2 bacteriophage RNA^23^, which is a homogeneous transcript of defined length. However, this model does not capture the heterogeneity inherent to cellular RNA. In the present study, we used total RNA extracted from *Escherichia coli* (*E. coli*), here referred to as e-RNAtotal, a complex and heterogeneous mixture of RNA species, to more accurately represent the physiological diversity of bacterial RNA. Using this RNA model, we demonstrate that MDP1 induces RNA condensation in a PTM-dependent manner, with the IDR playing a central role. These findings expand the known function of MDP1, showing that it not only compacts DNA but also contributes to RNA organization during dormancy. The identification of PTM-dependent RNA condensation opens new possibilities for targeting bacterial chromatin-like assemblies in persistent TB infections.

We first present the predicted domain organization of MDP1 using AlphaFold^24^ (Figure 1A). The structure reveals N-terminal HUR and C-terminal IDR, consistent with the classification of MDP1 as a histone-like protein^12,14,16–19,25–27^. Mass spectroscopic analyses have previously shown that the lysine residues of MDP1, particularly within its IDR, undergo extensive PTMs, mainly monomethylation along with dimethylation, trimethylation, acetylation and succinylation^13,16,26^ (Figures 1B and S1). Such PTMs are known in other intrinsically disordered proteins to modulate nucleic acid interactions by altering electrostatic charge, multivalency, and interaction specificity^28,29^. Particularly, acetylation and succinylation of MDP1 have also been reported to modulate its DNA-binding properties in Mtb^16,26,30^. These observations suggest that the IDR, along with its modification state, may similarly regulate RNA–MDP1 interactions. These analyses suggest that the IDR along with its modification state, may similarly regulate the RNA-MDP1 interaction.

**Figure 1.**
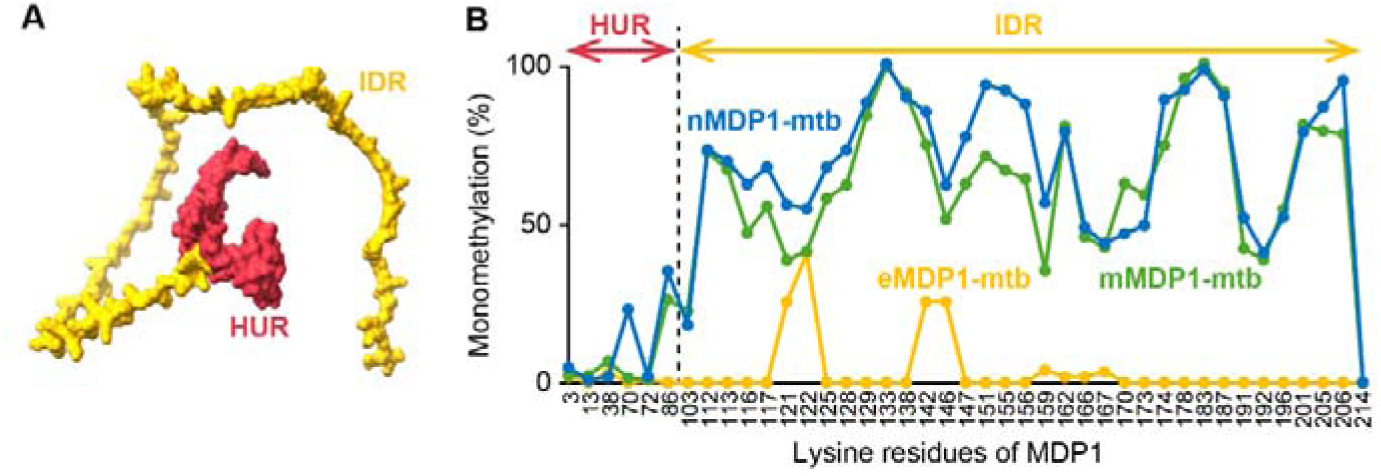
Predicted structure and PTM profile of MDP1. (A) Predicted 3D structure of MDP1 generated by AlphaFold. Structured HUR and string-like IDR are shown in red and yellow colors, respectively. (B) Relative abundance of monomethylation on lysine residues identified by mass spectrometry. Line plots of nMDP1-mtb, mMDP1-mtb and eMDP1-mtb are shown in blue, green and yellow colors, respectively^4,13^. Also see Figure S1 for the profiles of dimethylation, trimethylation, acetylation and succinylation.

To assess whether IDR and PTMs of MDP1 contribute to RNA interaction, we used e-RNAtotal as a model substrate. Two full-length MDP1 variants were tested: native MDP1 purified from Mtb (nMDP1-mtb), which carries native PTMs, and recombinant MDP1 expressed in *E. coli* (eMDP1-mtb), which lacks these modifications^13^.

First, we performed OM imaging to visualize RNA-protein interactions. Incubation of e-RNAtotal with nMDP1-mtb resulted in the formation of numerous globular condensates which increased in number and occupied a progressively larger proportion of the observed area from ∼30 to ∼60 min. These condensates initially appeared more globular at ∼30 min but further merging resulted in a less globular appearance (Figure 2Ai-ii). No such structures were formed when either RNA or proteins were incubated alone, under identical experimental conditions (Figure S2). Quantitative image analysis showed that condensates area increased over time, from ∼3% at ∼30 min to ∼6% at ∼60 min (Figure 2B), while circularity remained high (∼0.85; Figure 2C), indicating stable globular morphology. The area of individual condensates predominantly clustered ∼1 μm^2^, with no objects exceeding 50 μm^2^ (Figures 2D,E and S3A).

**Figure 2.**
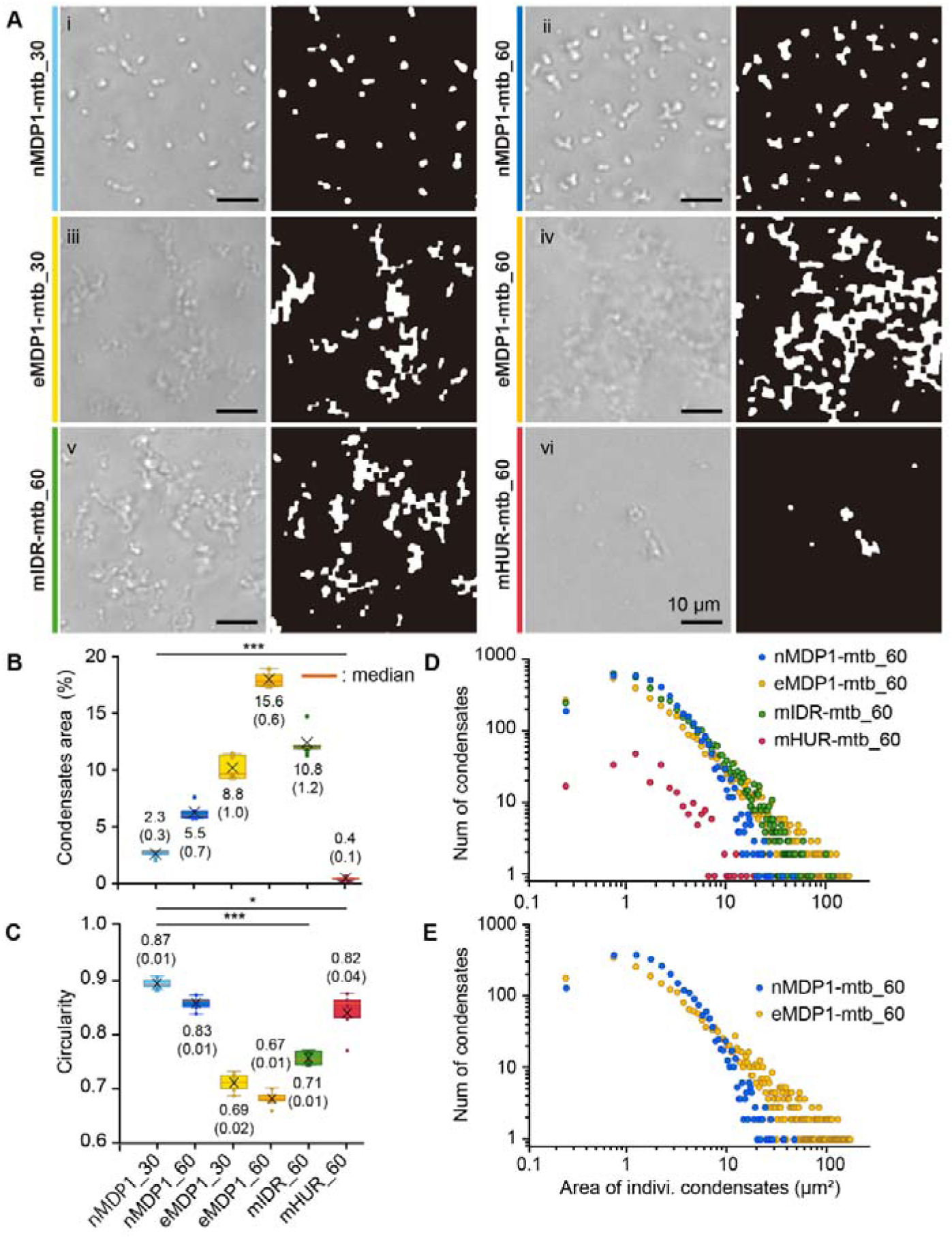
OM shows MDP1 induces RNA condensation in a variant-dependent manner. (A) Left (both sets of images), phase-contrast images taken at ∼30 or ∼60 min after mixing e-RNAtotal and indicated MDP1 variants. Right (both sets of images), corresponding binary images for quantification (see Methods). **(i–ii)** nMDP1-mtb, **(iii– iv)** eMDP1-mtb, **(v)** mIDR-mtb and **(vi)** mHUR-mtb. Time points of incubation are denoted by _30 or _60 on protein labels, scale bar, 10 μm. (B-E) Quantification of RNA-protein interactions based on three independent experiments. (B) Percentage area covered by condensates compared to total observation area. (C) Circularity of condensates. Statistics values are indicated as mean ± SD (SD in brackets) from three replicates. p < 0.05 was considered significant, \**p* < 0.05, \*\*\**p* < 0.001, *p* values are based on two sample *t*-test test. (D) Distribution of the area of individual condensates for all observations at ∼60 min. (E) Comparison between nMDP1-mtb and eMDP1-mtb is shown separately, extracted from (D).

In contrast, eMDP1-mtb induced formation of irregular, interconnected RNA condensates resembling chain-like networks (Figure 2Aiii–iv). These condensates covered larger proportion of the area (∼10% at ∼30 min and ∼18% at ∼60 min; Figure 2B) and exhibited lower circularity of ∼0.70 (Figure 2C). Distribution of the area of individual condensates included larger structures of >100 μm². It is important to note that these condensates appeared low-contrast under OM, potentially leading to an underestimation of their true size.

For dissecting the domain-specific contribution in RNA condensation, we next examined the MDP1 truncations originating from Mtb and expressed in *Mycobacterium smegmatis* (Msm). We were unable to generate these mutants in Mtb because of the essential role of MDP1 in bacterial growth, which makes it challenging to construct Mtb strains expressing only the HUR or IDR domains. The PTM status of these isolated constructs has not yet been characterized, but it is presumed to be lower than that of native MDP1 (Figure 1B).

The isolated IDR (mIDR-mtb) formed irregular, moderately sized condensates with RNA (Figure 2Av), covering ∼12% of the total observed area, exhibiting intermediate circularity of ∼0.75 (Figure 2B,C) and a broad distribution of the area of individual condensates of 1–100μm² (Figures 2D and S3C,D). The contrast of these condensates was intermediate between that of nMDP1-mtb and eMDP1-mtb, suggesting a moderate fusion efficiency. The HUR domain alone (mHUR-mtb), when incubated with RNA, produced rare, small clusters of ∼1–20μm² with high circularity of ∼0.85 but minimal area coverage of <1% (Figures 2Avi, B–D and S3D). This RNA-interacting ability of HUR is consistent with that of bacterial HU proteins, which bind to RNA, utilizing flexible β-ribbon arms extending from their head domain^31^.

These OM findings suggest that RNA condensation depends on both the degree of PTMs (Figure 2Ai-iv) and the critical role of the IDR (Figure 2Av-vi). While OM provided valuable insight into the overall morphology and distribution of RNA condensates, the limited resolution of this technique made it difficult to assess the fine structural details and nanoscale dynamics. To directly visualize these interactions at high spatial and temporal resolution, we next employed HS-AFM.

Control imaging of e-RNAtotal without protein revealed heterogeneous, extended features with multiple bends and branches distributed across the mica surface (Figure S4A and Movie S1). These structures were digested upon RNase treatment, confirming that these are RNA indeed (Figure S5 and Movie S2). Morphological analysis of e-RNAtotal over a ∼60 sec time window showed the maximum height range of ∼1.0–2.5 nm and the median volumes up to ∼1000 nm^3^ with circularity of ∼0.15 (Figures S4B,C and 4; NP), reflecting loosely folded single RNA strands lying flat on the mica surface. These parameters establish a baseline for subsequent analyses of RNA-protein interactions.

Upon addition of nMDP1-mtb, we observed rapid and robust binding of the protein to RNA, followed by the progressive formation of compact, globular condensates. These condensates continued to develop between ∼20 and ∼60 min (Figure 3A and Movie S3). This growth was driven by frequent fusion events between individual condensates (Figure 3E and Movie S4). Quantitative analysis indicated that the maximum height of these condensates increased from ∼7nm by ∼20 min to ∼10 nm by ∼40–60 min. Also, their volume increased largely, with a median of ∼21,000nm³ and some exceeding 40,000nm³ by ∼60 min (Figure 4A, upper panel). In parallel, the median circularity of these condensates increased to ∼0.7 by ∼60 min, aligning with globular morphology.

**Figure 3.**
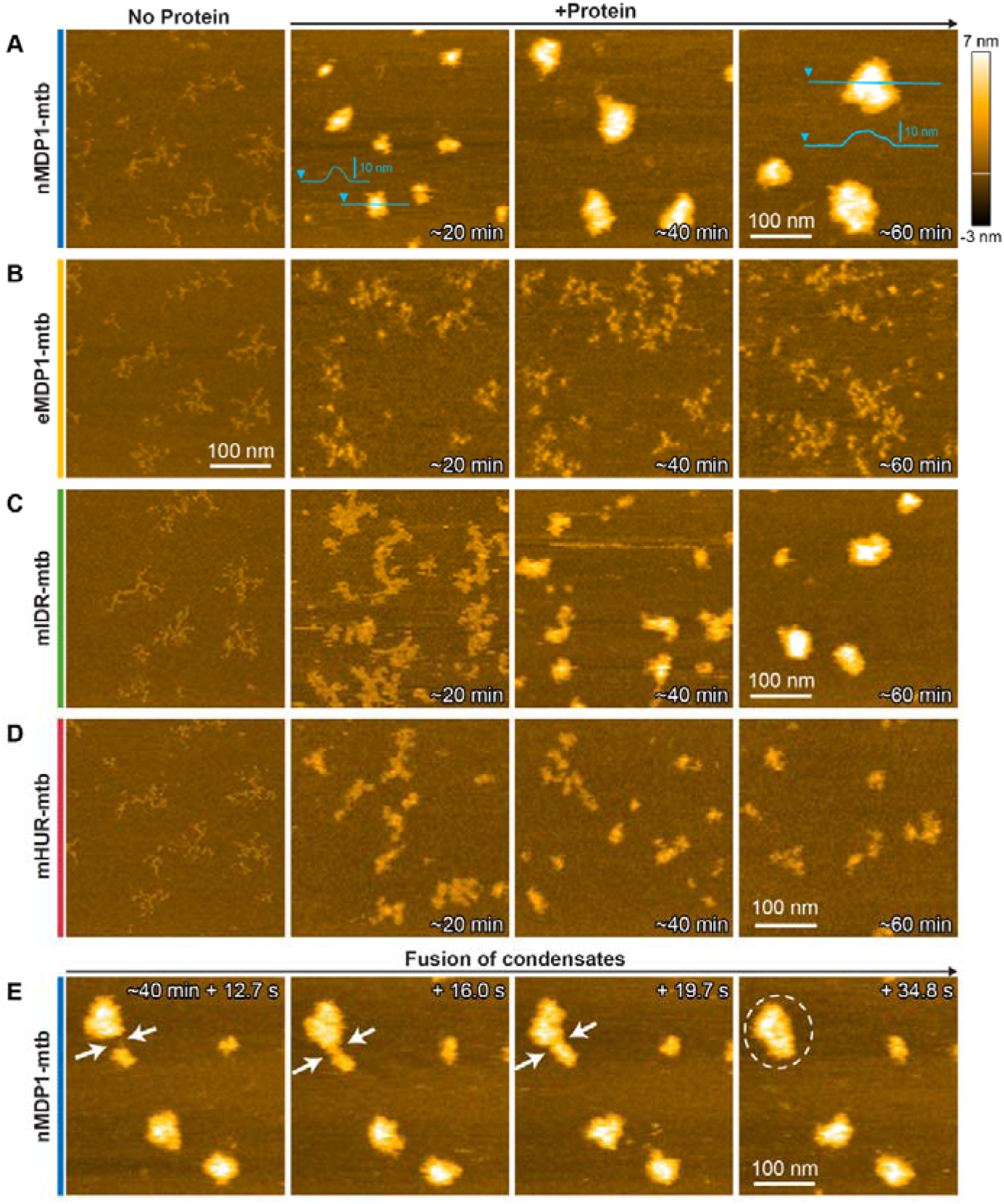
HS-AFM captures nanoscopic dynamics of RNA condensation induced by MDP1 variants. (A-D) Time lapse HS-AFM images taken before and after the addition of (A) nMDP1-mtb (Movie S3), (B) eMDP1-mtb (Movie S5), (C) mIDR-mtb (Movie S6), and (D) mHUR-mtb (Movie S7), at ∼20, ∼40, and ∼60 min. (E) Fusion process of nMDP1-mtb-induced condensates observed ∼40 min post incubation (Movie S4). White arrows indicate the gap and fusion sites while dashed circle shows the fused condensates. Contrasts of the images are normalized from-3 nm to 7 nm for direct comparison, white horizontal line in the color scale bar indicates “0 nm”. Height profiles are shown in light blue color in (A). High-contrast images for eMDP1-mtb and mHUR-mtb are shown in Figure S6. All experiments were conducted within a 350nm × 350nm scanning area with 100 × 100 pixels. Frame rates are shown in corresponding movie captions.

**Figure 4.**
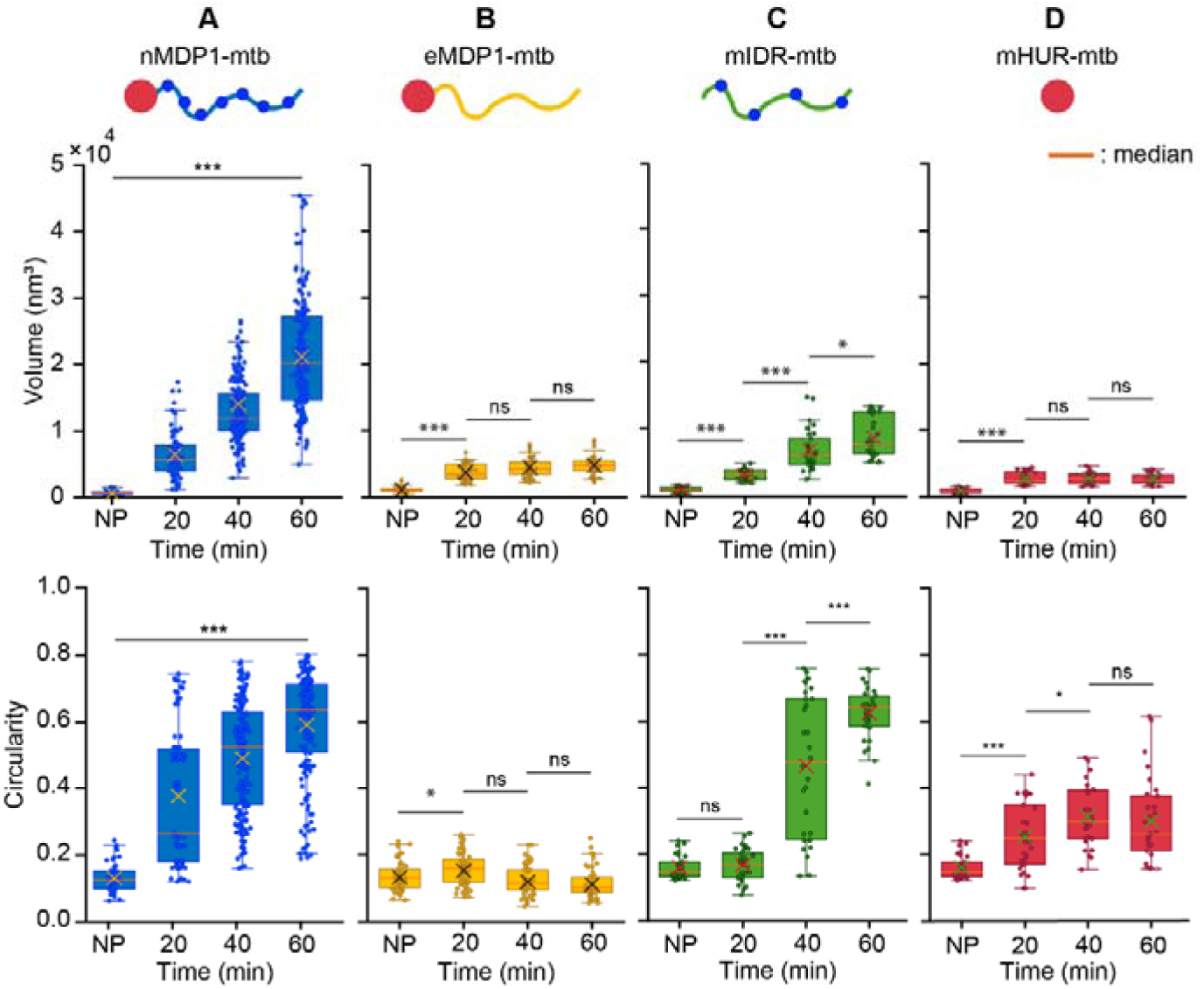
Quantitative HS-AFM analysis reveals PTM-enriched IDR and HUR cooperatively induce RNA condensation. (A-D) The top row shows volume distribution of individual control RNA and RNA-protein complexes, bottom row indicates the circularity. (A) nMDP1-mtb, (B) eMDP1-mtb, (C) mIDR-mtb, and (D) mHUR-mtb. Box plots display the distribution of volume for RNA only (NP; no protein) and after the addition of the protein, across indicated time points. The boxes represent the interquartile range. The orange horizontal line within each box marks the median, and the cross (×) indicates the mean. Whiskers extend to the most extreme data points within 1.5×IQR (inter-quartile range) from the quartiles; data points outside this range are considered outliers. Statistical comparisons were performed using two-sample *t*-test, one-way ANOVA, or Kruskal–Wallis test, followed by Tukey’s or Dunn’s post hoc test, depending on data distribution. *p* ≤ 0.05 was considered significant; ns = not significant, \**p* < 0.05, \*\*\**p* < 0.001.

These structural changes are consistent with the dynamics observed at the micron scale under OM.

In comparison, when RNA was incubated with eMDP1-mtb, numerous small globules were observed on RNA molecules (Figure S6A and Movie S8), leading to the formation of RNA-protein complexes. These complexes exhibited transient and unstable connections (Figure S6B), resulting into loose associations that were notably weaker than those observed with nMDP1-mtb. Consequently, RNA–protein complexes remained dispersed, or they formed irregular, chain-like clusters throughout the observation period (Figure 3B and Movie S5). Only a slight increase in height (∼4nm by ∼20 min) was detected with no further changes (Figure 3B), and also the overall morphology of the RNA–protein complexes showed little to no change. Throughout the ∼60 min incubation, the median volume and circularity remained low, ∼4,000 nm³ and ∼0.1, respectively (Figure 4B). This observation is consistent with that of OM in terms of the failure of PTM-deprived protein to induce globular RNA condensates.

When we incubated RNA with mIDR-mtb, no globules were observed on RNA molecules. Immediately after protein addition, the conformation of RNA began to change as the protein bound to it, but initially the complexes remained in an extended morphology. Although these extended complexes interacted with each other by ∼20 min, transformation into globular condensates was only observed after ∼40 min (Figure 3C and Movie S6). These condensates resembled those seen with nMDP1-mtb, but their growth was delayed under identical experimental conditions. The maximum height of these condensates increased from ∼4nm by ∼20 min to ∼10 nm by ∼60 min. The volume and circularity measurements showed a distinct pattern (Figure 4C). Around 20 min, volume increased significantly with a slight increase in circularity. However, a much sharper increase in both parameters was observed by ∼40 min. This sudden change reflects a transformation of mIDR-mtb–RNA complexes into globular condensates, resulting in a wider distribution of volume and circularity values. These condensates further fused together, resulting in increased compaction. By ∼60 min, the condensates reached a median volume of ∼8,000nm³ and a circularity of ∼0.6 (Figure 4C). These results indicate less-efficient condensation as compared to the full-length nMDP1-mtb, presumably due to the absence of HUR domain and lower level of PTMs.

When we observed the interaction of mHUR-mtb with RNA, small globules formed on RNA which caused local changes in RNA conformation (Figures 3D, S6C,D, Movie S7 and S8). However, the individual complexes rarely interact with each other. Quantification analysis showed an early increase in maximum height, reaching ∼4nm (Figure 3D), which then remained unchanged. During the first ∼20 min, median volume and circularity also showed moderate increase, reaching up to ∼3,000nm³ and ∼0.3 (Figure 4D), respectively, with no significant changes observed thereafter. These subtle changes suggest local RNA remodeling without full condensation.

Although both OM and HS-AFM consistently support the cooperation of IDR and HUR and the role of PTMs in RNA condensation, notable differences in condensate morphology and size trends were observed due to the distinct nature of each technique (Figure 5). For example, HS-AFM provides the nanoscopic view of individual molecules in action while it restricts the free diffusion because of surface adsorption of the sample. For eMDP1-mtb, HS-AFM showed loosely connected RNA-protein complexes (Figures 3B and S6A), but they failed to fuse extensively due to surface adsorption (Figure S6B). While in OM, free diffusion allowed these complexes to interact easily, resulting in their extremely large size (Figure 2). However, this protein could not induce efficient RNA condensation under both experimental setups (Figure 5B).

**Figure 5.**
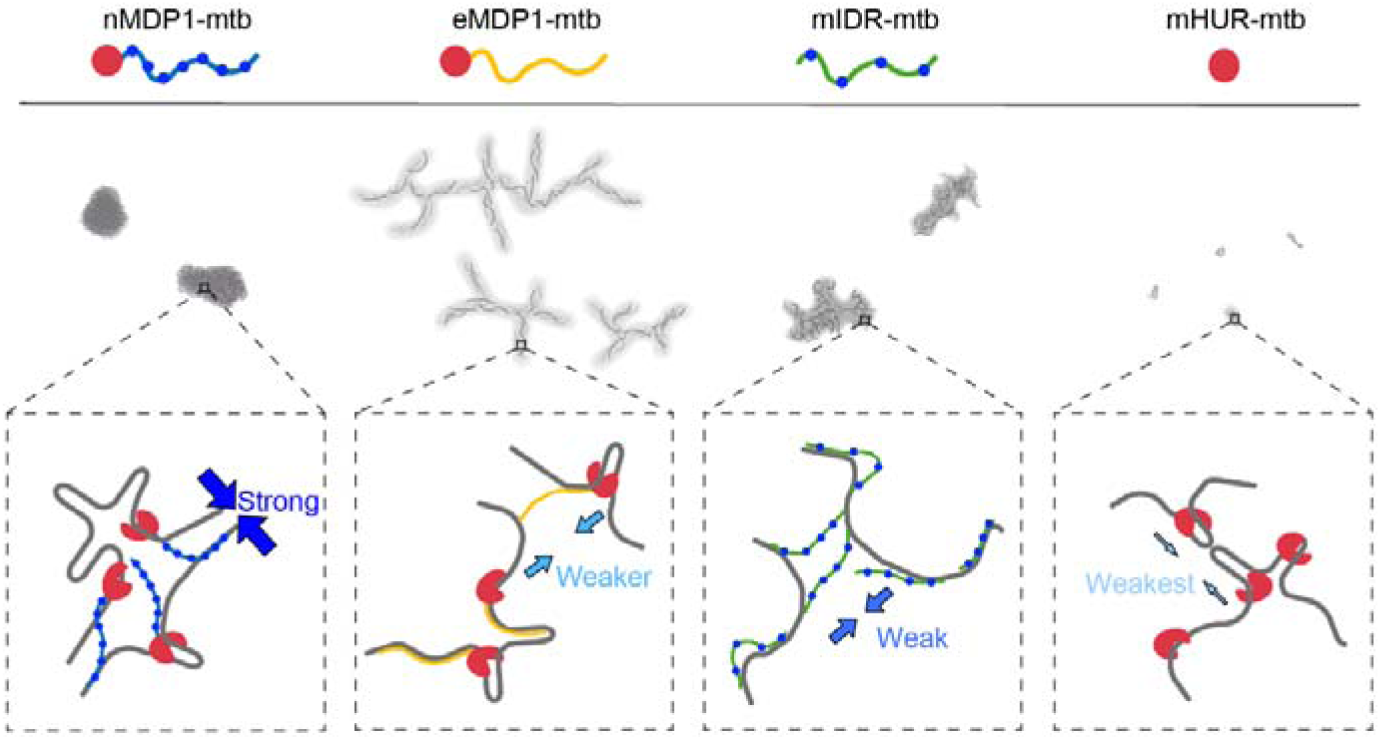
Model illustrating RNA condensation by MDP1 variants. Top: Cartoon representations of RNA interactions with indicated proteins, based on OM observations. Bottom: Schematic illustrations of RNA–protein interactions at the nanoscale, inferred from HS-AFM. Grey lines, RNA molecules; grey fillings, the shape of condensates while colour intensity from dark to light grey reflects the strength of condensation; arrows, relative RNA-binding strength and extent of fusion between RNA–protein complexes.

In case of mIDR-mtb, globular-like condensates were observed under HS-AFM similar to that of nMDP1-mtb (Figure 3A,C), but theses condensates appeared chain-like under OM (Figure 2). In HS-AFM, we observed considerable delay in growth of these condensates (Figure 3C), indicating weaker interaction ability. Conversely, in OM, these complexes could interact efficiently, due to free diffusion, like eMDP1-mtb, and therefore appeared chain-like (Figure 5C). This notion is further supported by the contrast observed in OM images, with the highest contrast for nMDP1-mtb, followed by mIDR-mtb and eMDP1-mtb while the size inversely follows this trend (Figure 2). Overall, the weaker fusion of RNA-protein complexes leads to reduced compaction power, resulting in larger sizes in solution.

When we observed the interaction of mHUR-mtb with RNA by HS-AFM, fusion of individual RNA-protein complexes was rarely observed (Figure 3D). Whereas, OM revealed micrometre-sized objects (Figure 2), although significantly smaller than those formed by other samples. This noticeable size increase visualized by OM, may be attributed to local conformational changes in RNA upon HUR binding^32^, which could expose additional RNA-binding sites and enable occasional interactions with complementary partners. Despite these interactions being infrequent, they may be sufficient to generate such assemblies (Figure 5D). The circularity of RNA-HUR complexes also differed between two techniques, with values ∼0.3 in HS-AFM and ∼0.85 in OM (Figures 2C and 4D). The high circularity observed in OM likely reflects early-stage RNA-protein binding events. Each complex appears round due to the limited resolution of OM, despite having a more irregular morphology at the nanoscale. In summary, these observations suggest that HUR-RNA complexes exhibit low inter-molecular binding efficiency, showing only weak and occasional interactions.

Taken together, we demonstrate that MDP1 induces RNA condensation *in vitro*, as visualized by HS-AFM and OM. This process depends on three key factors: the intrinsic role of the IDR, the synergy between the IDR and HUR domain, and the influence of PTMs. These findings introduce a novel aspect of RNA regulation in mycobacteria and expand the nucleic acid-binding function of MDP1 beyond DNA.

Our previous work demonstrated the binding of MDP1 with homogeneous MS2 RNA, having secondary structure^23^. In the present study, using a heterogeneous RNA substrate, we observed RNA condensation, an activity distinct from nonspecific aggregation. Our observations reveal that full-length nMDP1-mtb, enriched with PTMs, efficiently induces RNA condensation (Figures 2-4). Conversely, eMDP1-mtb, which lacks these modifications, shows markedly reduced condensation activity despite possessing both IDR and HUR (Figures 2-4). These results highlight the critical role of PTMs in modulating RNA-binding and condensation capability of MDP1, possibly due to their influence on electrostatic or conformational properties^33–35^. While well-studied in eukaryotic RNA-binding proteins^36,37^, such PTM-mediated regulation remains relatively underexplored in bacterial systems^38^.

Further, while the IDR alone does not induce efficient RNA condensation (Figures 2-4), the HUR domain contributes to RNA remodeling possibly by exposing potential interaction sites and promoting weak inter-RNA interactions, though significant fusion events are rare (Figures 2-4 and S6C-D). These observations highlight the complementary roles of HUR domain in initiating RNA remodeling, as reported for DNA^32^ and the PTM-enriched IDR in facilitating multivalent interactions necessary for effective condensation^39,40^. Together, these domains exhibit a cooperative dynamic essential for RNA condensation ability of MDP1^18,32,41^.

From a physiological perspective, MDP1-induced RNA condensation may play a crucial role in mycobacterial stress responses, particularly during dormancy, when both transcription and translation are downregulated globally^2,8,42,43^. Condensation may either stabilize transcripts by limiting their accessibility to ribosomes and RNases, or facilitate their degradation, thereby contributing to post-transcriptional transcriptome reorganization^44,45^. This RNA remodeling mirrors the established role of MDP1 organizing DNA into protective nucleoid structures^3,46^, further supporting its multifunctionality in nucleic acid organization. The dependence on PTMs suggests a dynamic regulatory mechanism, allowing MDP1 to respond to environmental cues and modulate gene expression beyond transcriptional control^4,13,16^.

While our findings highlight the role of MDP1 in RNA condensation, it is important to note the limitations of our study. The *in vitro* setting and the use of *E. coli* RNA may not fully reflect the intracellular environment or RNA composition relevant to mycobacterial physiology. Future research should explore how RNA length and sequence influence MDP1-induced condensation. It will also be important to identify the specific PTMs involved, uncover their regulatory pathways, and assess whether similar condensation occurs *in vivo*, particularly during dormancy.

## ASSOCIATED CONTENT

### Supporting Information

Supporting Information is available free of charge.

Materials and methods, data analysis, supporting images and analysis, and supporting movie captions (PDF).

HS-AFM movies for RNA only, RNA digestion, and RNA interaction with tested proteins (MP4).

## Author Contributions

N.S., A.N., S.M., and N.K. designed the research; N.S., A.N., K.N., Y.G., P.D.N.N., S.M., and N.K. performed the research; S.M., and A.N. prepared the protein samples; N.S., K.N., Y.G., H.I., K.U., and N.K. analyzed the data; N.S., and N.K. wrote the first draft, and the final draft was prepared after receiving input from all authors.

## Notes

The authors declare no competing financial interest.

## Supporting information

Supporting Information for main text

## ACKNOWLEDGMENT

We thank Prof. Toshio Ando, Ms. Kayo Nakatani and Ms. Risa Omura at Kanazawa University for the technical support of HS-AFM experiments, and Ms. Kaede Takada at Niigata University for technical support of bacterial strain construction.

## BRIEFS

Native MDP1 induces efficient RNA condensation.

## SYNOPSIS

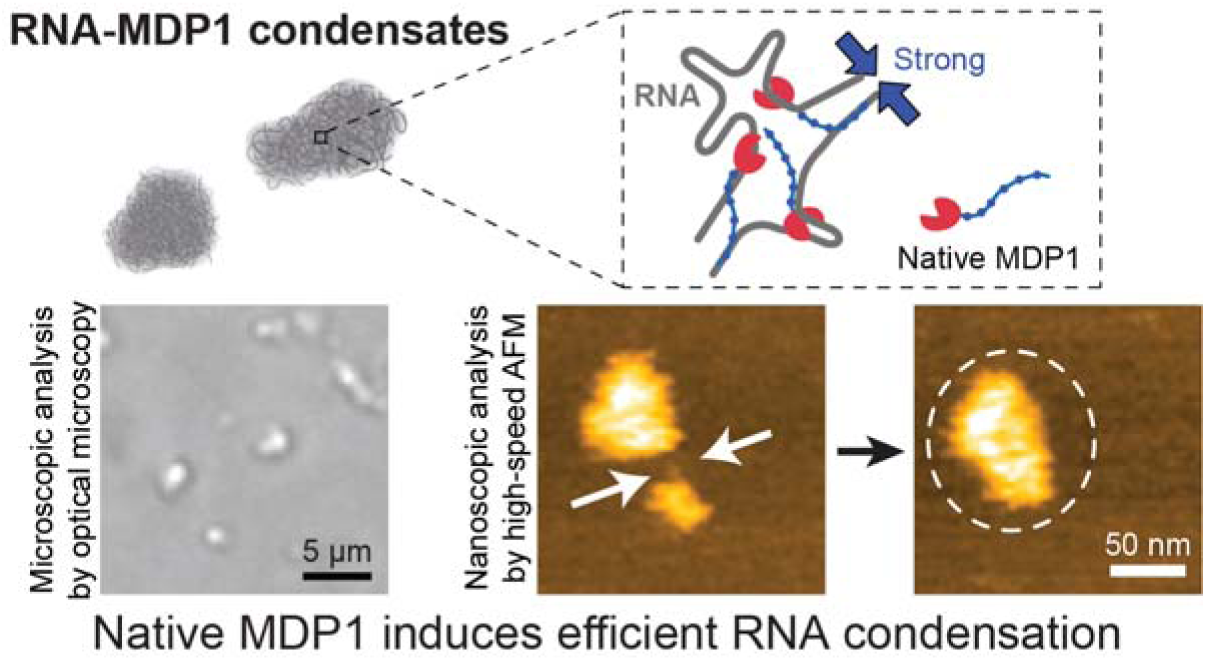

