## Supporting Information for main text for "Mycobacterial DNA-binding protein 1 (MDP1) induces RNA condensation as revealed by high-speed AFM"

**This PDF file includes:**

Materials and methods

Figures S1 to S6

Captions for Supporting movies 1 to 8

**Other Supporting materials for this manuscript include the following:**

Supporting movies 1 to 8

### Materials and methods

#### 1. RNA and RNase

Total RNA from *E. coli* (e-RNAtotal) isolated from DH5α cells, was purchased from ThermoFisher (Cat No. AM7000) and RNase (Ribonuclease A from Bovine Pancreas) was purchased from NACALAI TESQUE, INC (Product No. 30142-75).

#### 2. Purification of MDP1 proteins

##### 2.1 Native MDP1 purification

Native MDP1 (nMDP1-mtb) was purified from *M. tuberculosis* based on the method previously described[1]. *M. tuberculosis* H37Rv was cultured in Sauton medium at 37°C. After harvesting the bacteria by filtration, the bacterial cells were ground in a mortar with a pestle using quartz sand as an abrasive. The resulting lysates were then soaked in 1.5 l of 0.25 N HCl at 4°C for two nights. Acid-soluble proteins were obtained by sequential filtration through 0.45 µm and then 0.22 µm membrane filters. The filtrates were subsequently neutralized to approximately pH 7 using NaOH. Next, the extract was purified using a cation exchange column (HiPrep CM FF 16/10) (Cytiva). Gradient elution based on increasing salt concentration was performed, and the fractions containing the target protein were collected. After concentrating the eluted fractions, further purification was carried out by gel filtration chromatography using a HiLoad 26/60 Superdex 200 pg column (Cytiva), and the fractions containing MDP1 were again collected. The collected samples were then concentrated and dialyzed against a solution containing 8 M urea in 50 mM phosphate buffer (pH 6.8) with 300 mM NaCl. Refolding was performed by gradually reducing the urea concentration to 4 M, 2 M, 1 M, 0.5 M, and finally 0 M. The refolded protein was then used for further experiments. Approximately 7.8 mg of nMDP1-mtb was purified from 1 kg of wet bacterial cells.

##### 2.2 Recombinant eMDP1-mtb preparation

Recombinant MDP1 (eMDP1-mtb) was prepared by expressing MDP1 plasmid without any tag by inserting a TAA stop codon immediately after the MDP1 coding sequence in the pSO-ACE-HupB (Rv2986c)-His plasmid, which expresses His-tag (6 histidine tag) MDP1 in *E. coli* (eMDP1-mtb)[2]. This modified plasmid, designated pET-22b-MDP1(op, no-tag), was introduced into *Rosetta2(DE3) pLysS*. The recombinant bacteria were cultured in LB medium using a jar fermenter. A soluble protein fraction was prepared from the cultured bacterial cells and purified using a cation exchange column (HiPrep CM). Buffer A consisted of 25 mM phosphate buffer (pH 6.8) containing 150 mM NaCl, while Buffer B consisted of 50 mM phosphate buffer (pH 6.8) containing 2 M NaCl. Stepwise elution was performed, and the target protein was eluted in the fraction corresponding to 35% Buffer B. The collected fraction was subsequently dialyzed against 50 mM phosphate buffer (pH 6.8) containing 300 mM NaCl. Gradient elution based on increasing salt concentration was then performed using Buffer B, resulting in the isolation of highly purified eMDP1-mtb protein. Protein refolding was carried out using urea, following the same procedure used for nMDP1-mtb, and the refolded protein was used for further observations. Approximately 1.4 mg of eMDP1-mtb was purified from 1 l of recombinant *E. coli* culture.

**2.3 Recombinant mIDR-mtb and mHUR-mtb preparation**

To prepare mIDR-mtb, sequences encoding His-tag (6 histidine tag) mIDR of $MDP1_{mtb}$ (mIDR-mtb, derived from Rv2986c of *Mtb* H37Rv strain) were amplified using previously constructed pSO246-ami-mdp1-mtb (a plasmid having *mdp1-mtb*[3], by PCR using the primer sets: 5'-GTCCATATGCCCGCTGTTAAGCGTGGTGT-3' and 5'-TTTGCAAGCAGCAGATTACG-3'. Amplified DNA fragments were digested with NdeI and KpnI (Takara Bio, Shiga, Japan) and then inserted between NdeI and KpnI sites at the downstream of inducible promoter/regulator system[4,5] (cloned from acetamidase gene locus [*ami*] of *Msm* $mc^2$_155 of pSOami[3] (kanamycin [Km] resistant [$Km^R$]). Obtained plasmid was finally designated as pSOami-mIDR-mtb. *Msm* Δ*mdp1* (hygromycin B [Hyg] resistant [$Hyg^R$], kindly provided by Dr. John L. Dahl, University of Minnesota Duluth)[6] was transformed with pSOami-mIDR-mtb and colonies harboring pSOami-mIDR-mtb were selected on Middlebrook 7H10 agar plates (BD) supplemented with 0.5% (v/v) glycerol, 10% OADC enrichment (final concentrations of the components in the agar plates: 0.5% [w/v] bovine serum albumin [FUJIFILM Wako Pure Chemical Corporation, Osaka, Japan], 0.081% [w/v] NaCl, 0.2% [w/v] D-glucose, and 0.006% [v/v] oleic acid), 50 μg/ml Hyg, and 10 μg/ml Km. Protein expression of the selected clones was examined in culture in the presence of 0.2% acetamide (Ace).

For preparation of mHUR-mtb, we used *Msm* Δ*mdp1* strains harboring the genes for His-tag HUR of MDP1-mtb (mHUR-mtb) as described previously[4]. As for purification of mHUR-mtb from *Msm*, a strain described above was inoculated in 200 ml of Mueller-Hinton II Broth, Cation-Adjusted, (MHB, BD) supplemented with 0.05% (v/v) Tween 80, 50 μg/ml Hyg, and 10 μg/ml Km (MHB/Tw80/Hyg/Km) and pre-cultured at 37°C with vertical shaking (120 rpm). At ~0.7 optical density at 600 nm ($OD_{600}$), 50 ml aliquots of pre-culture were added to 450 ml MHB/Tw80/Hyg/Km supplemented with 2% Ace (final 2%) and incubated for 48 h. Bacterial cells were harvested, washed, and stored at -80°C until use. After 48 h culture (mHUR-mtb induction), bacterial cells were harvested by centrifugation at 8,000 rpm (10,590 ×g), 4°C, for 10 min, and then washed once with ultra-pure water. Bacterial cell pellets were then stored at -80℃ until use.

For purification of mIDR-mtb, frozen stocks of the strains described above were inoculated in MHB/Tw80/Hyg/Km and cultured at 37°C with vertical shaking (120 rpm). At around 1.0 $OD_{600}$, Ace (final 2%) was added to the culture and incubated again for 36 h. Bacterial cells were harvested, washed, and stored at -80°C until use.

Proteins were purified from the frozen bacterial pellets as described previously[3]. Purity of the protein was determined by SDS-PAGE. Fractions containing His-tagged proteins were combined and dialyzed with phosphate-buffered saline (PBS[-]) for subsequent experiments. Purified proteins were then concentrated using Amicon Ultra-15 Centrifugal Filter with appropriate pore size (Merck Millipore, Burlington, MA). Protein concentrations were determined using Pierce™ BCA Protein Assay Kit (ThermoFisher Scientific, Waltham, MA).

#### 3. Optical Microscopy (OM)

Optical microscopy experiments were performed to visualize MDP1-mediated RNA condensation. e-RNAtotal was used at a final concentration of 54 ng/µl. Prior to imaging, RNA and protein samples were diluted in phosphate-buffered saline (PBS). For each observation, 4 µl of RNA solution and 4 µl of protein solution (final protein concentration: 5 µM) were gently mixed directly in individual wells of a CORNING 384-well imaging plate (Cat No. 3540, Corning, Corning, NY). The samples were imaged using a phase-contrast optical microscope (BZ-X10, Keyence, Japan), at room temperature (23-25°C). Phase-contrast images were captured using a 20× objective lens combined with 3× digital zoom. Each well was monitored continuously for a total duration of ~60 min. For nMDP1-mtb and eMDP1-mtb, around 15 snapshots were taken at ~30 min and ~60 min after mixing, while for mIDR-mtb, mHUR-mtb and controls, snapshots were recorded after ~60 min of incubation only. Each experiment was performed in triplicate.

#### 3.1 Data analysis of OM images

For image quantification, 5 representative images were used, although >15 images were taken for each condition. These images of TIFF-format with a resolution of 1,920 × 1,440 pixels corresponding to 240 µm × 180 µm were analysed using ImageJ[7] to evaluate condensates properties following background and noise removal. First, Gaussian Blur (σ = 3) was applied to reduce noise and smooth particle contours. Background subtraction was then performed using the rolling ball algorithm (radius = 20 pixels) to enhance contrast between particles and the background. Automatic thresholding was used to binarize the images, minimizing misidentification or subjective interpretation. The images were masked, eroded 2 times, dilated 6 times, followed by 4 times erosion to refine particle boundaries.

Then, we used the command of "analysed particles" to obtain number, area, and circularity of individual condensates. The percentage area covered by condensates compared to total observation area was calculated by summing up areas of individual condensates and then dividing by whole imaging area (240 µm × 180 µm), for one image. These calculations were repeated for 5 different images per condition, and then the corresponding box-and-whisker plot was generated. To assess circularity, average circularities of individual condensates were calculated for one image. The box-and-whisker plot was generated similarly using 5 different images per condition. The area distribution of individual condensates was analysed using all obtained areas of individual condensates from 5 images, per condition.

**4. Observation of RNA-protein complexes by HS-AFM**

HS-AFM imaging in tapping mode was performed in solution at room temperature (24-26°C) using a custom-built HS-AFM, as described previously[3]. We used small cantilever (BL-AC10DS-A2, Olympus; spring constant, ~0.1 N/m; resonant frequency and quality factor in liquid, ~500 kHz and ~1.5, respectively). The free oscillation amplitude ($A_0$) was set to 1.5–2.5 nm, and the set-point amplitude was maintained at ~90% of $A_0$ for optimal tip–sample interaction. A small cylindrical glass stage (2 mm diameter × 2 mm height, Japan Cell) was affixed to the top of a Z-scanner using nail polish, and a thin muscovite mica disc (1.5 mm diameter, 0.05 mm thickness, Furuuchi Chemical Corporation) was glued onto the stage with epoxy.

To visualize RNA, 2 µl e-RNAtotal (final concentration ~5 nM) was diluted in scanning buffer (10 mM Tris-HCl, pH 7.5, 20 mM NaCl, 60 mM KCl), deposited onto the freshly cleaved mica surface, and incubated for 3 min to allow partial adsorption. The surface was gently rinsed with the same scanning buffer (20 µl) to remove unbound molecules, and imaging was conducted in the same buffer (60 µl). To ensure that the observed molecules are RNA, RNase was added to the imaging chamber at final concentration of ~1 nM, and the digestion process was observed. For time-lapse RNA-protein interaction experiments, HS-AFM imaging was initiated after RNA adsorption, and proteins (nMDP1-mtb, eMDP1-mtb, mIDR-mtb and mHUR-mtb) were added directly to the imaging chamber at final concentrations of ~200 nM. Morphological changes in RNA and RNA–protein complexes were recorded in real time. All experiments were conducted more than three times, except for the interactions of mIDR-mtb and mHUR-mtb with RNA, which were performed twice. The representative images and movies for each experimental condition in this study originate from a single experiment, showing sequential imaging of the observed events.

#### 4.1 Data analysis of HS-AFM images

We analyzed HS-AFM images using UMEX, a custom-built software. As a pre-treatment of analysis, we applied a low-pass filter to eliminate spike noise and a flattening filter to level the xy-plane. The maximum height of each RNA molecule was measured by calculating the difference between the maximum height of RNA, and the average height of the substrate. For volume and circularity measurements, the outlines of individual molecules were manually traced, and these parameters were calculated.

All statistical analyses were performed using OriginPro 2024 (OriginLab Corporation, USA). A significance cutoff of $p = 0.05$ was used for all tests. The normality of distributions was assessed using the Shapiro–Wilk and Kolmogorov–Smirnov tests. For data that followed a normal distribution (mIDR-mtb), comparisons were conducted using two-sample *t*-test or one-way ANOVA, followed by Tukey’s post hoc test where applicable. For data not normally distributed (nMDP1-mtb, eMDP1-mtb, and mHUR-mtb), the non-parametric Kruskal–Wallis test was applied, followed by Dunn’s post hoc test to determine group-wise differences between the control (no protein) and experimental conditions (after protein addition).

**Supplementary Figures**

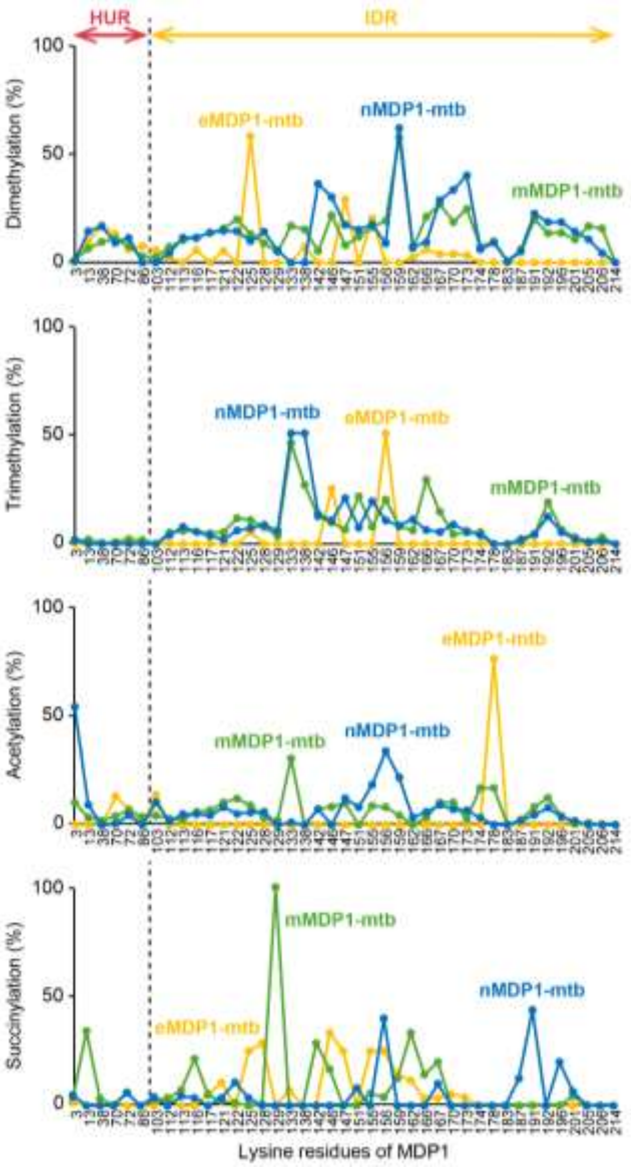


**Figure S1. Extended PTM profile of MDP1.**

Relative abundance of dimethylation, trimethylation, acetylation and succinylation on lysine residues identified by mass spectrometry. Line plots of nMDP1-mtb, mMDP1-mtb and eMDP1-mtb are shown in blue, green and yellow colors, respectively[8,9].

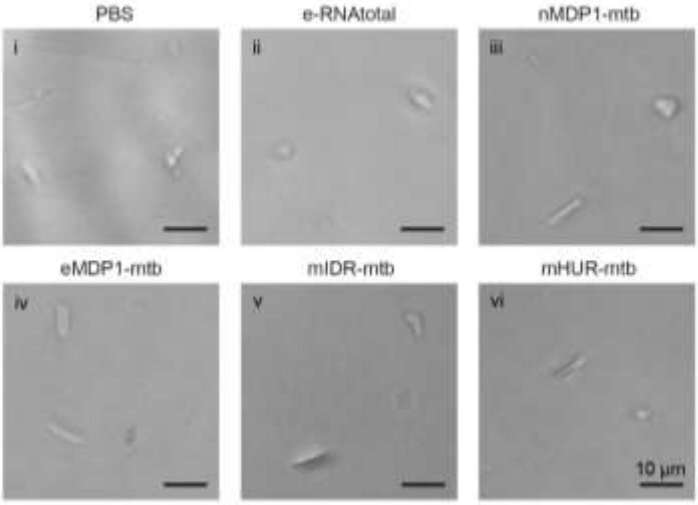


**Figure S2. OM imaging of control samples.**

OM images showing control conditions, in which the indicated RNA or protein samples were observed individually: (i) PBS alone, (ii) e-RNAtotal, (iii) nMDP1-mtb, (iv) eMDP1-mtb, (v) mIDR-mtb, and (vi) mHUR-mtb. Scale bars: 10 µm. No condensation or aggregation structures as shown in Figure 2A were observed under these control conditions. The particles visible in the images are likely impurities or dust and are shown intentionally to demonstrate that the imaging conditions were suitable to detect aggregates or condensates, if formed.

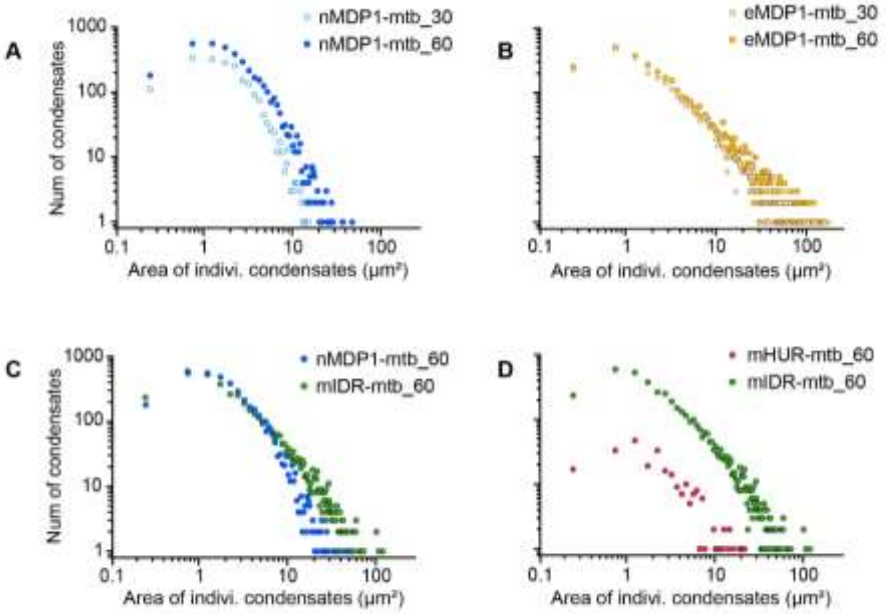


**Figure S3. Extended quantitative analysis of OM images.**

Distribution of the area of individual condensates for individual observations at indicated time points. (A) nMDP1-mtb at ~30 (blue outlined circle) and ~60 min (blue circle) of incubation. (B) eMDP1-mtb at ~30 (yellow outlined circle) and ~60 min (yellow circle) of incubation. (C) nMDP1-mtb (blue circle) and mIDR-mtb (green circle) at ~60 min of incubation. (D) mHUR-mtb (red circles) in comparison with mIDR-mtb (green circles) at ~60 min of incubation.

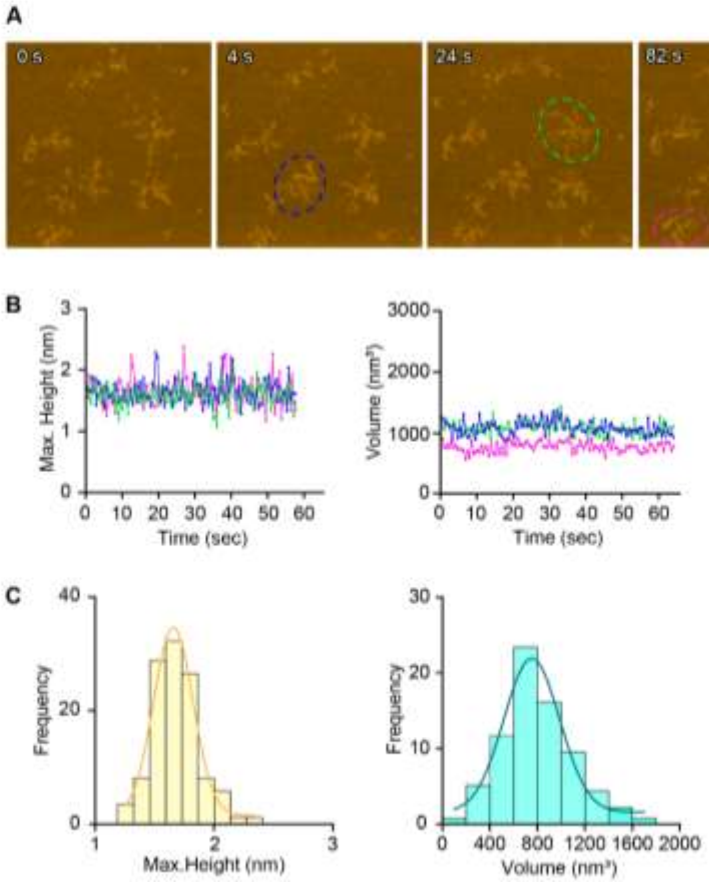


**Figure S4. Dynamics of e-RNAtotal observed by HS-AFM.**

(A) Time-lapse HS-AFM imaging of e-RNAtotal. Three representative RNA molecules (blue, green, magenta; dashed ovals) tracked over ~60 sec, images taken from Movie S1, scale bar: 100 nm. Contrasts of the images are normalized from -3 nm to 7 nm for direct comparison, white horizontal line in the color scale bar indicates “0 nm”. (B) Quantitative tracking of individual RNA molecules showing temporal changes in height (left), and volume (right). (C) Distribution of RNA maximum height and volume. RNA height was determined by the maximum height of each molecule respective to the substrate.

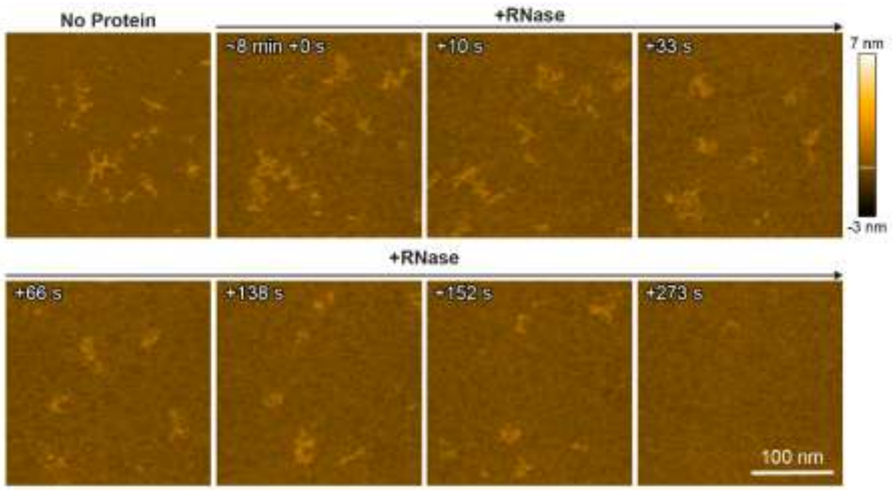


**Figure S5. e-RNAtotal digestion by RNase observed by HS-AFM.**

Time-lapse HS-AFM imaging of e-RNAtotal digestion by RNase. Contrasts of the images are normalized from -3 nm to 7 nm for direct comparison, white horizontal line in the color scale bar indicates “0 nm”. This experiment was conducted within a 250 nm × 250 nm scanning area with 100 × 100 pixels. Images taken from Movie S2, scale bar: 100 nm.

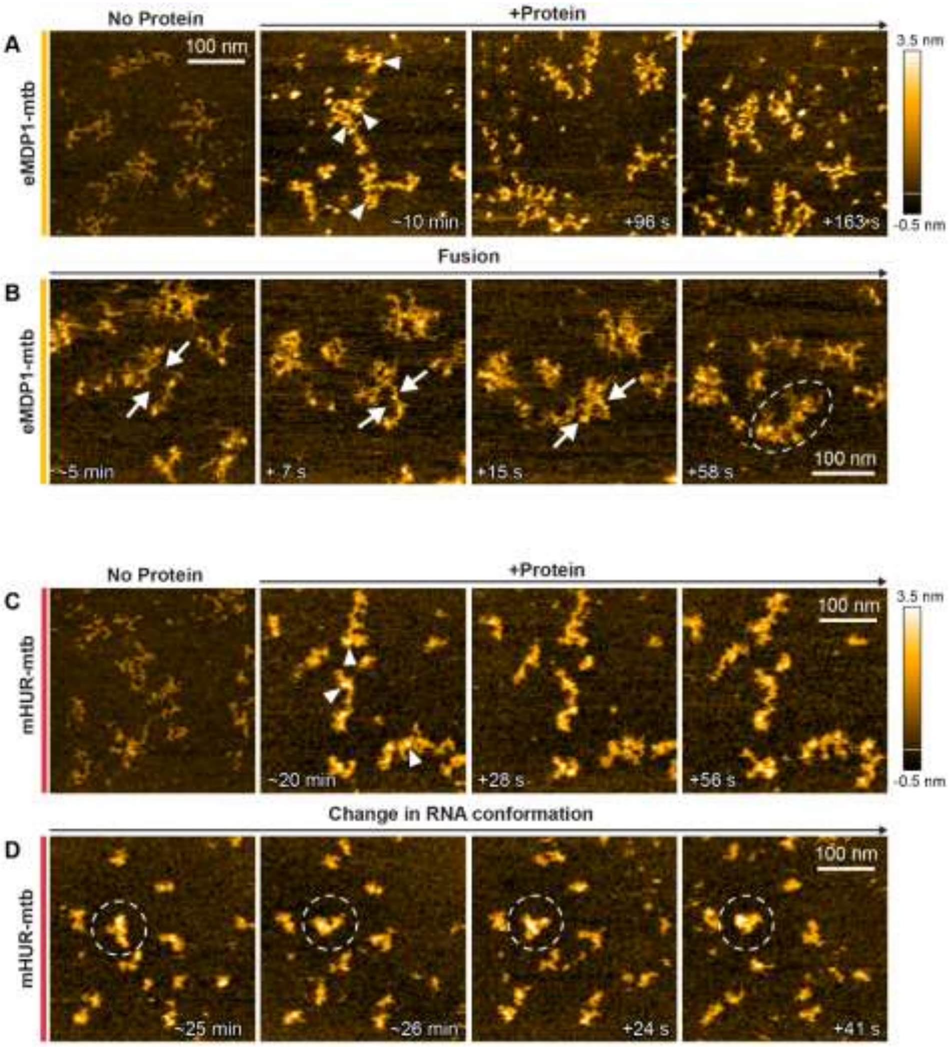


**Figure S6. High-contrast nanoscopic view of eMDP1-mtb and mHUR-mtb binding to e-RNAtotal.**

(A) Representative images showing RNA morphology before and after the addition of eMDP1-mtb, white arrowheads indicate typical protein molecules bound to RNA. (B) Time-lapse images showing the progressive fusion of eMDP1-mtb-bound RNAs after ~5 min of protein addition (scanning area 300 nm × 300 nm with 150 × 150 pixels). White arrows indicate merging RNA-protein complexes, and the final frame highlights a fused structure (dashed oval). (C) Representative images showing RNA morphology before and after the addition of mHUR-mtb, white arrowheads indicate the binding of typical proteins on RNA. (D) Time-lapse images of mHUR-mtb–bound RNA showing gradual changes in RNA conformation after ~25 min of protein addition. Dashed circles highlight selected RNA regions undergoing conformational rearrangement. Contrasts of the images are normalized from -0.5 nm to 3.5 nm for better visualization, white horizontal line in the color scale bar indicates “0 nm”, scale bars, 100 nm.

**Supporting movies captions**

**Movie S1:** Time-lapse HS-AFM movie showing e-RNAtotal molecules. Representative frames are presented in Figure S4. Scanning area is 350 nm × 350 nm with 100 × 100 pixels. The imaging rate was ~2.44 frames per second (fps), and the movie is played at ~10 fps. The contrast of the movie is normalized from -3 nm to 7 nm: scale bar, 100 nm.

**Movie S2:** Time-lapse HS-AFM movie showing e-RNAtotal digestion by RNase at indicated time points. Representative frames are presented in Figure S5. Scanning area is 250 nm × 250 nm with 100 × 100 pixels; imaging rate was ~2.44 fps, and the movie is played at ~30 fps. The contrast of the movie is normalized from -3 nm to 7 nm: scale bar, 100 nm.

**Movie S3:** Time-lapse HS-AFM movie showing e-RNAtotal only (No protein) and RNA-protein interaction after the addition of nMDP1-mtb at ~20, ~40 and ~60 min; white arrow indicates the addition of the protein. Representative frames are presented in Figure 3A. Scanning area is 350 nm × 350 nm with100 × 100 pixels. The imaging rates were ~2.44 fps for 0 min (no protein), and ~2 fps, ~2.44 fps and ~1.33 fps for ~20, ~40 and ~60 min, and the corresponding movie panels are played at ~10 fps,~8 fps,~10 fps, and ~6 fps, respectively. The contrast of the movie is normalized from -3 nm to 7 nm; scale bar: 100 nm.

**Movie S4:** Time-lapse HS-AFM movie showing the fusion process of nMDP1-mtb-induced condensates observed ~40 min after the addition of the protein. White arrows indicate the gap and fusion sites while dashed circle shows the fused condensates. Representative frames are presented in Figure 3E. Scanning area is 350 nm × 350 nm with 100 × 100 pixels. The imaging rate was ~2.44 fps and the movie is played ~10 fps. The contrast of the movie is normalized from -3 nm to 7 nm; scale bar: 100 nm.

**Movie S5:** Time-lapse HS-AFM movie showing e-RNAtotal only (No protein) and RNA-protein interaction after the addition of eMDP1-mtb at ~20, ~40 and ~60 min; white arrow indicates the addition of the protein. Representative frames are presented in Figure 3B. Scanning area is 350 nm × 350 nm with 100 × 100 pixels. The imaging rates were ~2.44 fps for 0 min and 1.44 fps for ~20, ~40 and ~60 min, and the corresponding movie panels are played at ~10 fps and ~6 fps, respectively. The contrast of the movie is normalized from -3 nm to 7 nm; scale bar: 100 nm.

**Movie S6:** Time-lapse HS-AFM movie showing e-RNAtotal only (No protein) and RNA-protein interaction after the addition of mIDR-mtb; white arrow indicates the addition of the protein. Representative frames are presented in Figure 3C. Scanning area is 350 nm × 350 nm with 100 × 100 pixels. The imaging rates were ~2.44 fps for 0 min and ~1.66 fps for ~20, ~40 and ~60 min, and the corresponding movie panels are played at ~10 fps and ~7 fps, respectively. The contrast of the movie is normalized from -3 nm to 7 nm; scale bar: 100 nm.

**Movie S7:** Time-lapse HS-AFM movie showing e-RNAtotal only (No protein) and RNA-protein interaction after the addition of mHUR-mtb; white arrow indicates the addition of the protein. Representative frames are presented in Figure 3D. Scanning area is 350 nm × 350 nm with 100 × 100 pixels. The imaging rates were ~2.44 fps for 0 min and ~1.66 fps for ~20, ~40 and ~60 min, and the corresponding movie panels are played at ~10 fps and ~7 fps, respectively. The contrast of the movie is normalized from -3 nm to 7 nm; scale bar: 100 nm.

**Movie S8:** Time-lapse high-contrast HS-AFM movie showing the interaction of eMDP1-mtb (left panel) and mHUR-mtb (right panel) with e-RNAtotal at indicated time points. Representative frames are presented in Figure S6A and S6C, respectively. Scanning area is 350 nm × 350 nm with 100 × 100 pixels. The imaging rates were ~1. 44 fps and ~1.66 fps for eMDP1-mtb and mHUR-mtb, and the corresponding movie panels are played at ~6 fps and ~7 fps, respectively. The contrast of the movie is normalized from -0.5 nm to 3.5 nm for better visualization; scale bar: 100 nm.
